# FOXM1 Modulation Alleviates Epithelial Remodeling and Inflammation in Eosinophilic Esophagitis

**DOI:** 10.1101/2025.05.25.655133

**Authors:** Masaru Sasaki, Joshua X. Wang, Yusen Zhou, Kanak V. Kennedy, Ryugo Teranishi, Takefumi Itami, Satoshi Ishikawa, Takeo Hara, Heidi Winters, Emily A. McMillan, Mark Mahon, Hailey Golden, Diya Dhakal, Alyssa Bacarella, Chizoba N. Umeweni, Benjamin J. Wilkins, Tatiana A. Karakasheva, Kelly A. Whelan, Sydney M. Shaffer, Melanie A. Ruffner, Amanda B. Muir

**Author notes:** Correspondence Amanda B. Muir, Perelman School of Medicine, University of Pennsylvania, Abramson Research Center 902E, 3615 Civic Center Boulevard, Philadelphia, PA 19104, USA. **Abbreviations** AKT, protein kinase B; ALI, air-liquid interface; ChIP, chromatin immunoprecipitation; DAPI, 4′,6-diamidino-2-phenylindole; DEG, differentially expressed gene; DMSO, dimethyl sulfoxide; EoE, eosinophilic esophagitis; GSEA, Gene Set Enrichment Analysis; H&E, hematoxylin and eosin; IL, interleukin; KSFM, keratinocyte-serum free medium; FOXM1, forkhead boxM1; OFR, organoid formation rate; OVA, ovalbumin; PDO, patient-derived organoid; PI3K, phosphatidylinositol 3-kinase; PID, Pathway Interaction Database; RCM-1, Robert Costa Memorial drug-1; RT-PCR, reverse transcription-polymerase chain reaction; TEER, transepithelial electrical resistance; Th2, T-helper type 2. Disclosures Amanda B. Muir has served on the medical advisory boards for Nexstone Immunology and Bristol Meyers Squib, Regeneron/Sanofi, and Uniquity. All of the other authors declare that they have no conflicts of interest. Author Contributions MS, TAK, and ABM were responsible for the study concept and design. HW, MS, JXW, KVK, RT, and TI performed the experiments. MS, JXW, YZ, KVK, RT, TI, SI, TH, AB, BJW, and ABM performed the data analyses. MS and NNU created the schematics. EAM provided reagents. HG, MM, and DD recruited patients. MS, TAK, and ABM wrote the original draft, and MS, TAK, KAW, SMS, MAR, and ABM edited it. All authors discussed the results and reviewed the manuscript.

## Abstract

**Background:** Eosinophilic esophagitis (EoE) is a chronic allergic disease characterized by esophageal epithelial remodeling, barrier dysfunction, and inflammation. Despite histologic remission, molecular and structural changes in the epithelium persist, contributing to ongoing symptoms and relapse. The transcription factor FOXM1 has been shown to be a key regulator of epithelial proliferation and inflammation in allergic asthma.

**Objective:** To investigate the role of FOXM1 in epithelial disruption in EoE and to evaluate the therapeutic potential of FOXM1 inhibition.

**Design:** FOXM1 expression was analyzed in human esophageal biopsies, patient-derived organoids, and murine EoE models. IL-13 stimulation was used to model EoE in vitro. The effects of FOXM1 inhibition via the small molecule RCM-1 and siRNA-mediated knockdown were assessed by histology, gene expression profiling, organoid formation rates, and barrier integrity assays. RNA sequencing and chromatin immunoprecipitation were performed to elucidate molecular mechanisms.

**Results:** FOXM1 was significantly upregulated in patients with active EoE and localized to the basal epithelium. IL-13 increased FOXM1 expression, which impaired epithelial differentiation and enhanced basal cell hyperplasia. FOXM1 inhibition restored differentiation markers, reduced basal hyperplasia, and improved barrier function. In murine models, RCM-1 ameliorated epithelial changes and decreased eosinophil infiltration. Mechanistically, FOXM1 directly regulated cell cycle gene, CCNB1, which was upregulated in EoE and downregulated upon FOXM1 inhibition. FOXM1 expression was driven by an IL-13-PI3K/AKT axis.

**Conclusion:** FOXM1 plays a pivotal role in epithelial disruption in EoE by driving proliferation and impairing differentiation. Targeting FOXM1 restores epithelial homeostasis, mitigates inflammation, and offers a novel therapeutic approach for EoE.

**Key Messages:** **What is already known on this topic:** Eosinophilic esophagitis is marked by epithelial remodeling and barrier dysfunction driven by Th2 inflammation. Despite remission, molecular and histologic changes in the esophageal epithelium persist, contributing to symptoms and relapse. The mechanisms underlying this epithelial dysregulation remain poorly understood.

**What this study adds:** This study identifies FOXM1 as a key transcriptional regulator of epithelial disruption in EoE, demonstrating that FOXM1 inhibition restores epithelial differentiation, reduces basal cell hyperplasia, improves barrier integrity, and mitigates inflammation.

**How this study might affect research, practice, or policy:** Targeting FOXM1 offers a novel therapeutic strategy to restore epithelial homeostasis and reduce inflammation in EoE. This dual approach, addressing both epithelial and immune dysregulation, may guide future therapeutic development and improve patient outcomes.

## Introduction

Eosinophilic esophagitis (EoE) is an allergic disease of the esophagus in which disruption of the esophageal mucosa and inflammation lead to vomiting, dysphagia, and eventually stricture^1^. Recent work has demonstrated that patients in remission, despite having a decreased inflammation, have ongoing molecular and histologic changes in the esophageal epithelium^2^. These changes contribute to ongoing symptomatology and allow for disease relapse^3,4^. There is an urgent need to define the mechanisms of epithelial dysregulation in order to achieve true long-lasting mucosal healing.

While the pathophysiology of EoE is incompletely understood, exposure to food and aeroallergens triggers a T-helper type 2 (Th2) cell production of interleukin-13 (IL-13). IL-13 in turn stimulates epithelial production of eotaxin-3, also known as C-C motif chemokine ligand 26 (CCL26), the major eosinophil chemoattractant^1,5,6^. Independent of its effects stimulating eosinophil chemotaxis, IL-13 has been shown in vitro and in vivo to trigger epithelial remodeling events including dilated intracellular spaces and basal cell hyperplasia^5–8^. Histologically, 95% of patients with active EoE, demonstrate hyperplasia of the basal epithelium^3^, which at the molecular level is a failure of cells to terminally differentiate, as shown by decreased expression of involucrin (IVL), filaggrin (FLG), and keratin-6^4,8,9^. Single-cell evaluation of the EoE epithelium has described an expansion of the proliferative progenitor population with retained Ki-67 and keratin-13. Understanding the underlying mechanism that contributes to this combination of increased proliferation and failure to differentiate of progenitor cells represents a knowledge gap in the field.

Restoring the proliferation and differentiation gradient in the EoE epithelium may lead to a durable healing and make the epithelium less prone to relapse. Forkhead box M1 (FOXM1), a transcription factor of the Forkhead box family, is recognized as a major regulator of cell cycle progression and the mitotic program^10,11^. In the lung, inhibition of FOXM1 prevents IL-13 mediated goblet cell hyperplasia in murine asthma models. Further, FOXM1 has recently been implicated in epithelial cell differentiation^12–15^, fibroblast activation^16^, and pathogenesis of allergen-induced inflammatory diseases^17,18^. Aberrant FOXM1 expression in the skin and oral cavity disrupted epithelial differentiation and homeostasis, resulting in hyperplasia^12,13^. In contrast, terminally differentiated skin keratinocytes attenuate FOXM1 expression and downregulation of FOXM1 enhances differentiation markers^14,15^, suggesting that FOXM1 is essential for the regulation of the epithelial proliferation-differentiation gradient.

In this study, we examined FOXM1 expression in the normal and EoE esophageal epithelium. We utilized patient-derived esophageal organoids and a murine model of EoE to examine the role of FOXM1 in driving epithelial disruption and propose this as a therapeutic target for allergic inflammation of the esophagus.

## Methods

### Monolayer culture of human esophageal cells

Immortalized human normal esophageal epithelial cells, EPC2-hTERT, were kept in keratinocyte-serum free medium with 0.09 mM Ca^2+^ (KSFM; Thermo Fisher Scientific, Waltham, MA). High-calcium (1.8 mM Ca^2+^) KSFM was used to induce differentiation^19^. Cells were stimulated by recombinant human IL-13, IL-4, tumor necrosis factor alpha (TNF-α), or transforming growth factor beta (TGF-β) to recapitulate EoE milieu. The Robert Costa Memorial drug-1 (RCM-1; S6898; Selleck Chemicals, Houston, TX) and LY294002 (S1105; Selleck Chemicals) was used to inhibit FOXM1 activity and phosphatidylinositol 3-kinase (PI3K) signaling, respectively^17,20^.

### Esophageal 3-dimensional (3D) organoid culture

EPC2-hTERT or primary patient-derived organoids (PDOs) were established as described previously^9,21^. Briefly, the prepared EPC2-hTERT cells and patient derived esophageal cells were suspended in Matrigel basement membrane matrix (Corning Inc., Corning, NY) and seeded under modified KSFM with 0.6 mM Ca^2+^. Organoids were cultured for 7 days and then treated with IL-13 (10 ng/ml), RCM-1 (10 μM), or vehicle (phosphate-buffered saline for IL-13 or dimethyl sulfoxide (DMSO) for RCM-1) for 4 days. The day 11 organoids were harvested for further analyses.

Organoid formation rate (OFR) was defined as the number of ≥50 μm organoids at day 11 divided by the total seeded cells in each well^7,9^.

### Air-liquid interface (ALI) culture system

EPC2-hTERT cells were cultured with the ALI system as described previously^7^. Cells were stimulated with IL-13 (10 ng/ml), RCM-1 (20 μM), or vehicle in the basolateral compartment from day 9 to 15. Transepithelial electrical resistance (TEER) was measured to evaluate epithelial barrier. The ALI-cultured epithelium at day 15 was harvested for RNA extraction and histology.

### Patients and esophageal specimens

After obtaining informed consent, distal esophageal specimens were derived from patients who underwent esophagogastroduodenoscopy. Based on diagnostic criteria of EoE^22^, we classified patients as active EoE, inactive EoE, and non-EoE. Patient demographics are shown in Supplemental Table S1. Patient and Public Involvement were not appropriate or possible to involve patients or the public in the design, or conduct, or reporting, or dissemination plans of our research.

### Murine EoE

EoE models were induced as described previously^21,23,24^. Brief schedule of experimental EoE mice is described in Figure 5A. Murine esophagi were harvested for flow cytometry (5 mice per group) and histology (10 hpf views randomly selected from 3 mice per group). All experiments were performed in accordance with the Institutional Animal Care and Use Committee of The Children’s Hospital of Philadelphia.

### RNA sequencing analysis on human esophageal biopsies

We utilized public RNA sequencing data (GSE58640)^25^ on esophageal biopsies derived from active EoE and control subjects. The raw data was processed as described previously^26^. Gene expression was generated by reads per kilobase per million (RPKM) normalization method. Differentially expressed genes (DEGs) were defined as fold change ≥1.5 or ≤2/3 and *P* <0.05. Enrichment pathway analysis was conducted via gene set enrichment analysis (GSEA; Broad Institute, Cambridge, MA) using genes with *P* <0.05^27^. Top 10 enriched pathways with the lowest false discovery rate (FDR) value in the Pathway Interaction Database (PID) in the EoE subjects were shown^28^.

### Quantitative reverse transcription-polymerase chain reaction (qRT-PCR)

RNA isolation and reverse transcription were performed as described previously^26,29^. The used TaqMan and SYBR Green primers are provided in Supplemental Table S2 and S3. Relative mRNA levels were standardized against *glyceraldehyde-3-phosphate dehydrogenase* (*GAPDH*) levels as a housekeeping control.

### Western Blot (WB)

Whole-cell lysates were extracted as described previously^7^. Primary antibodies used are provided in Supplemental Table S4. Clarity Western ECL Substrate (Bio-Rad, Hercules, CA) were used to detect immunoblots. β-actin was set as a loading control.

### Immunohistochemistry and immunofluorescence

Paraffin-embedded human and murine esophagi, organoids, and ALI-cultured epithelia were sectioned and subjected to hematoxylin and eosin (H&E) staining, immunohistochemistry, or immunofluorescence as described previously^7,9^. Primary antibodies used are provided in Supplemental Table S4. Stained slides were imaged with an All-in-One Fluorescence Microscope BZ-X710 (KEYENCE Corp., Osaka, Japan).

### Flow cytometry

Murine esophagi were dissociated and filtered in the same manner as human specimens described above. Eosinophils were identified as CD45^+^/CD11b^+^/Siglec F^+^/SSC^high^ cells^30^. BD LSRFortessa Cell Analyzer (BD Biosciences, Franklin Lakes, NJ) and FlowJo software (FlowJo LLC, Ashland, OR) were used for analyses.

### Small interfering RNA (siRNA) transfection

Silencer Select Pre-Designed siRNA against FOXM1 (s5250 or s5248; Life Technologies) or Silencer Select Negative Control No. 1 siRNA (4390844; Invitrogen, Waltham, MA) were transfected into cells at a final concentration of 20 nM using Lipofectamine RNAiMAX Transfection Reagent (13778150; Invitrogen) according to the manufacturer’s protocol.

### Gene expression analysis on siFOXM1-transfected cells

The day after siRNA transfection (siNC, siFOXM1_1, or siFOXM1_3), EPC2-hTERT cells were cultured with the high-calcium KSFM for 3 days in monolayer culture and then harvested for RNA sequencing. Library preparation, RNA sequencing, and alignment were performed as described previously^7^. Mapped reads were analyzed by DESeq2^31^. One sample from siNC cells was removed from subsequent analyses due to suspected contamination of other materials. Genes for which adjusted *P* values could not be calculated were excluded. DEGs were defined as fold change ≥1.5 or ≤2/3 and *P* <0.05. Using the overlapping DEGs in siFOXM1_1 and siFOXM1_3 cells, Gene Ontology analysis was carried out with DAVID bioinformatics software 6.8^32^. Top 5 enriched and depleted terms with the lowest FDR values in the siFOXM1 cells were indicated.

### Proliferation assay

The day after transfection, cells were reseeded with or without IL-13 (10 ng/ml) in a 96-well plate. Proliferative ability was measured by WST-1 reagent (Roche Holding AG, Basel, Switzerland) according to the manufacturer’s protocol. Cells were incubated with the reagent for 90 min at 37°C and absorbance was read at 450 nm with a reference reading at 600 nm.

### Cell cycle assay

Cells stained with propidium iodide were evaluated through BD LSRII (BD Biosciences). Cell cycle phases were provided by the Watson Pragmatic algorithm in FlowJo software.

### Chromatin Immunoprecipitation (ChIP) assay

ChIP samples were prepared with truChIP Chromatin Shearing Kit (520154; Covaris, Woburn, MA) and immunoprecipitation was performed with SimpleChIP Plus Sonication Chromatin IP Kit (Cell Signaling Technology, Danvers, MA) based on the manufacture’s protocol. Five μg of FOXM1 antibody (sc-376471 X; Santa Cruz Biotechnology, Dallas, TX) or IgG (2729; Cell Signaling Technology) was added and incubated. Finally, DNA was purified using the PCR purification kit and quantified by real-time qPCR. ChIP results were calculated by fold enrichment method against IgG. The primer against cyclin B1 (CCNB1) for ChIP-qPCR is provided in Supplemental Table S3.

### Statistical analysis

Data are indicated as mean ± standard deviation (SD). Variables were analyzed by two-tailed Student’s t-test for two-group comparison or one-way analysis of variance (ANOVA) for multigroup. Pearson correlation coefficient was used to measure linear association between two variables. Statistical analyses were performed with GraphPad Prism (GraphPad Software, San Diego, CA). A *P* value <0.05 was considered statistically significant.

Detailed materials and methods are provided in online supplemental files.

## Results

### FOXM1 expression is elevated in patients with active EoE and localizes to the basal epithelium

Utilizing previously published RNA sequencing from EoE patients and non-EoE controls^25^, we performed a Pathway Interaction Analysis. Multiple cell cycle-related pathways were enriched (Figure 1A). In the case of FOXM1, not only the pathway was upregulated in EoE patients, but the *FOXM1* gene itself was found to be upregulated in these patients (Figure 1B and C). We validated this finding by qRT-PCR, demonstrating increased *FOXM1* expression in active EoE patients compared to those in remission and non-EoE controls (Figure 1D). In longitudinal samples from patients who achieved histologic remission, *FOXM1* expression decreased between the active state and remission (Figure 1E). When biopsy specimens were stained for FOXM1, its expression was localized to the basal epithelium for all specimens, but in patients with active EoE it was notably increased (Figure 1F and G).

**Figure 1.**
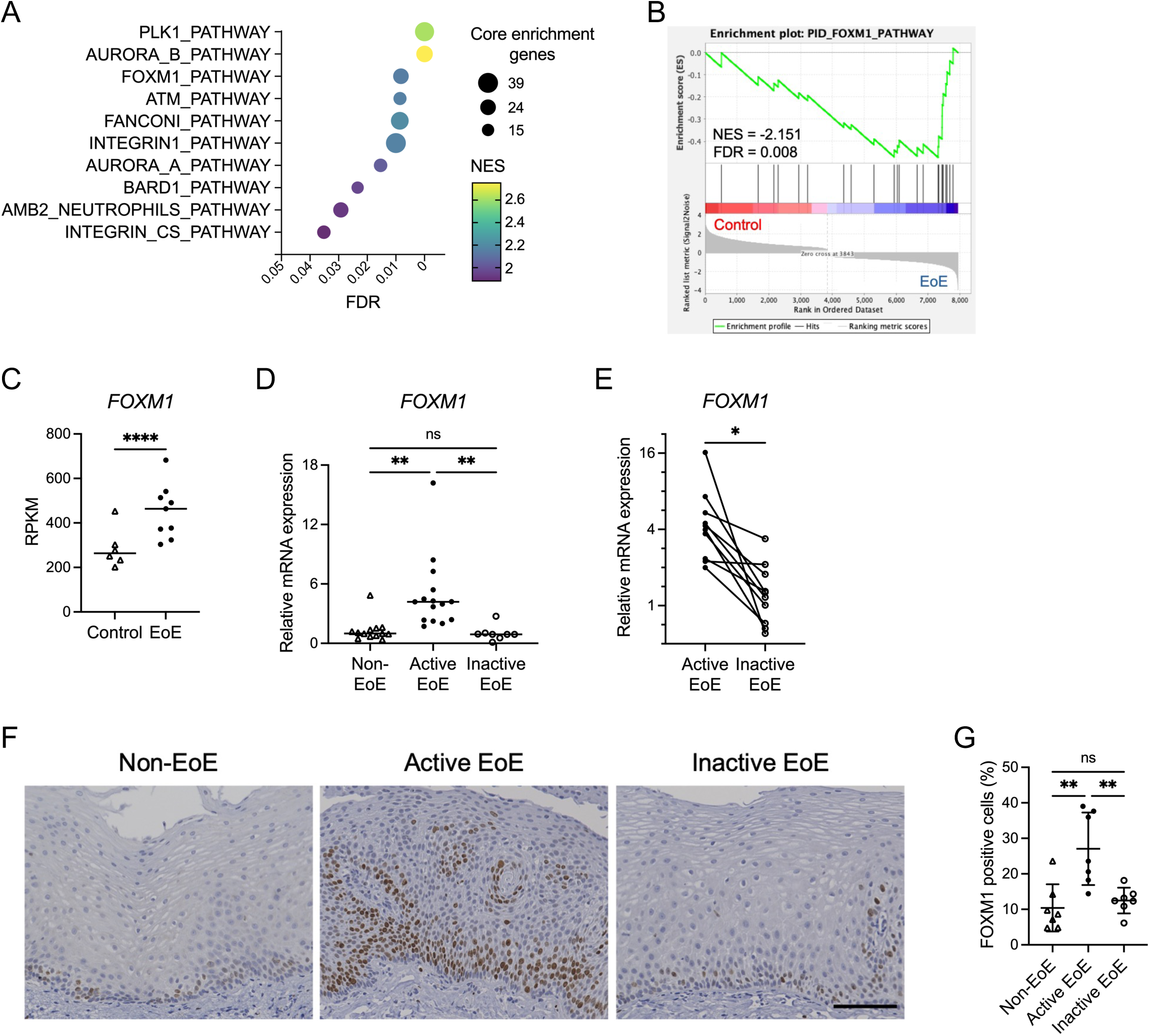
FOXM1 expression is elevated in patients with active EoE. (A-C) RNA sequencing analyses on the esophageal biopsy specimens (GSE58640). (A) Top 10 enriched pathways in Pathway Interaction Database (PID) in EoE patients generated by Gene Set Enrichment Analysis. (B) Enrichment plot of FOXM1 pathway in the PID. (C) Reads per kilobase per million (RPKM) values for FOXM1 in control (n = 6) and active EoE (n = 9) subjects. (D) Quantitative RT-PCR for *FOXM1* in non-EoE (n = 13), active EoE (n = 15), and inactive EoE (n = 8) biopsies in our cohort. Expressions are shown as relative values against the median expression in non-EoE group. (E) Changes in *FOXM1* mRNA expressions in active and inactive phases of the same patient (n = 10). (F and G) Representative images of immunohistochemistry for FOXM1 of non-EoE, active EoE, and inactive EoE biopsies. Scale bar, 100 μm. In the epithelium per high-power field, FOXM1 levels were quantified as number of nucleus-stained cells with FOXM1 divided by the total number of nuclei (n = 7 per group). Data are indicated as means ± SDs. Two-tailed Student’s t-test (C), one-way analysis of variance (D and G), and paired t-test (E) were utilized for statistics. \**P* <0.05, \*\**P* <0.01, \*\*\*\**P* <0.0001. FDR, false discovery rate; NES, normalized enrichment score; ns, not significant

### Th2 cytokines increase FOXM1 expression in the esophageal epithelium

In order to determine the factors driving FOXM1 expression in EoE, we stimulated EPC2-hTERT cells with the EoE relevant cytokines. We found that treatment with IL-13 and IL-4 (and not TNF-α or TGF-β) induced increased *FOXM1* expression in 2-dimensional culture (Figure 2A and B). As IL-13 stimulated cultures demonstrated the robust *FOXM1* expression and this is the major effector cytokine in EoE, we used IL-13 throughout this investigation. IL-13 stimulation of esophageal organoids (both EPC2-hTERT and PDO) demonstrated increased *FOXM1* expression by qRT-PCR (Figure 2C). Similar to patient biopsies, immunohistochemical staining of organoids showed localization of FOXM1 in the basal epithelium, which was increased with IL-13 stimulation (Figure 2D).

**Figure 2.**
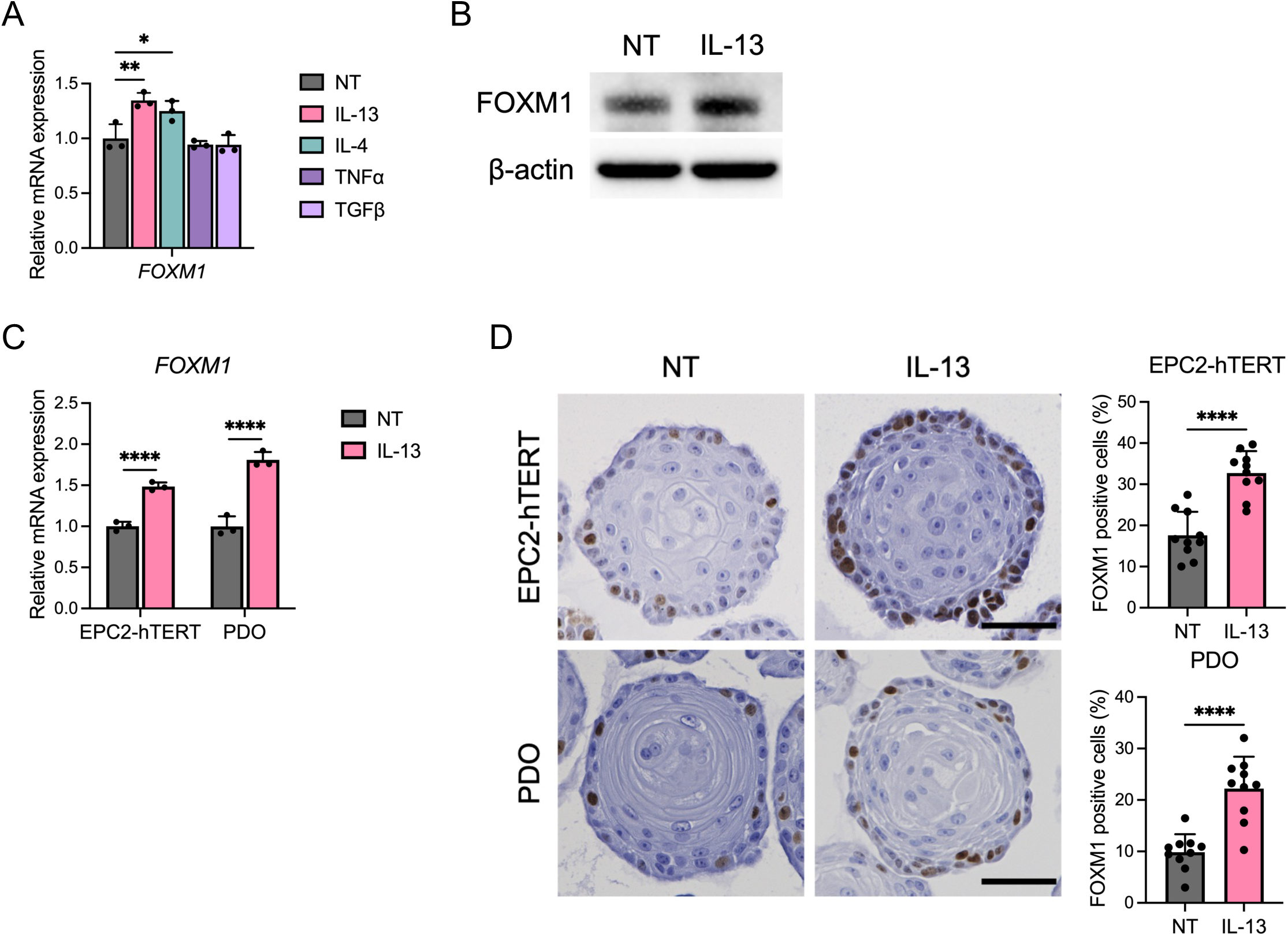
FOXM1 is upregulated by IL-13 stimulation in esophageal epithelium. (A) Quantitative RT-PCR for *FOXM1* of EPC2-hTERT cells stimulated with or without IL-13 (10 ng/ml), IL-4 (10 ng/ml), TNF-α (10 ng/ml), or TGF-β (10 ng/ml) for 24 h in monolayer culture (n = 3). (B) Representative images of immunoblot for FOXM1 of EPC2-hTERT cells stimulated with or without IL-13 (10 ng/ml) for 24 h in monolayer culture. (C and D) EPC2-hTERT or patient derived organoids (PDOs) were cultured for 7 days and then stimulated with or without IL-13 (10 ng/ml) for 4 days. The day 11 organoids were analyzed. (C) Quantitative RT-PCR for *FOXM1* of the organoids (n = 3). (D) Representative images of immunohistochemistry for FOXM1 of the organoids. Scale bar, 50 μm. Ratio of nucleus-stained cells with FOXM1 per organoid were counted (n = 10). Data are representative of three independent experiments and indicated as means ± SDs. One-way analysis of variance (A) and two-tailed Student’s t-test (C and D) were utilized for statistics. \**P* <0.05, \*\**P* <0.01, \*\*\*\**P* <0.0001. NT, nontreated

### Inhibition of FOXM1 attenuates IL-13-induced epithelial changes

Hyperproliferation of the basal epithelium and failure of cells to appropriately differentiate are hallmarks of EoE. To examine the role of FOXM1 on this proliferation-differentiation axis, we identified two silencing RNAs (siFOXM1_1 and siFOXM1_3) that reduced FOXM1 transcript and protein compared to scrambled control (siNC, Supplemental Figure S1A and B). Expression of the differentiation marker IVL was increased after FOXM1 knockdown, while expression of basal marker tumor protein p63 (TP63) was decreased after knockdown (Supplemental Figure S1C) even in the setting of IL-13 stimulation. These findings lead to the hypothesis that FOXM1 is a key regulator of the decision point between proliferation and differentiation in esophageal epithelium biology.

Next we utilized RCM-1, a small molecule which leads to proteasomal degradation of FOXM1^17^. We confirmed the dose-dependent loss of FOXM1 in epithelial cells treated with RCM-1 at 10 μM and 20 μM (Figure 3A). RCM-1 treatment enhanced epithelial differentiation as demonstrated by increased expression of IVL by WB, qRT-PCR, and staining in EPC2-hTERT cells and PDO cultures (Figure 3B-F and Supplemental Figure S2A-C). We further evaluated differentiation marker FLG with similar findings (Figure 3C-F). Thus, RCM-1 treatment rescued IL-13-induced reduction of these differentiation markers. While IL-13 treatment increased expression of basal cell markers TP63 and SRY-box transcription factor 2 (SOX2) and promoted basal cell hyperplasia, RCM-1 treatment suppressed this phenotype (Figure 3B-F and Supplemental Figure S2A-C).

**Figure 3.**
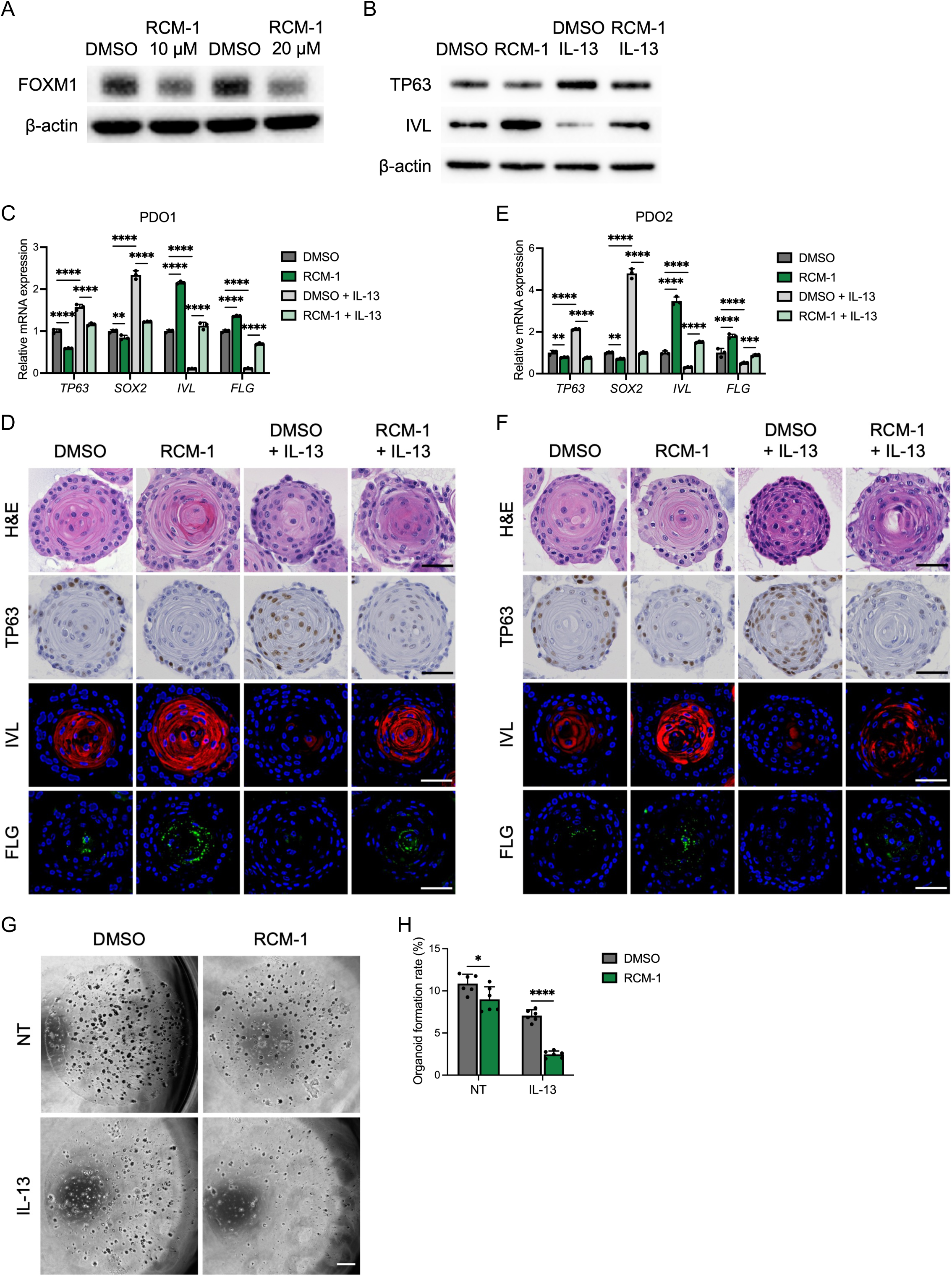
FOXM1 inhibition with RCM-1 abrogates epithelial disruption in the setting of IL-13 stimulation. (A) Representative images of immunoblot for FOXM1. Monolayer-cultured EPC2-hTERT cells were treated with DMSO or RCM-1 (10 μM or 20 μM) for 24 h. (B) Representative images of immunoblot for TP63 and IVL. EPC2-hTERT cells were treated with or without IL-13 (10 ng/ml) and RCM-1 (20 μM) in the setting of high-calcium KSFM (1.8 mM Ca^2+^). (C-F) Patient derived organoids (PDO1: C and D; PDO2: E and F) were cultured for 7 days and then treated with or without IL-13 (10 ng/ml) and RCM-1 (10 μM) for 4 days. Day 11 organoids were subjected to quantitative RT-PCR (n = 3), hematoxylin and eosin (H&E) staining, immunohistochemistry for TP63, and immunofluorescence staining for IVL (red) and FLG (green) of the organoids. Representative images are shown. DAPI (blue). Scale bar, 50 μm. (G and H) Representative phase contrast images of EPC2-hTERT organoids. Organoids were treated with or without IL-13 (10 ng/ml) and RCM-1 (10 μM) from day 7 to day 11 and then passaged. Organoid formation rate (OFR) was assessed at day 11 (passage 1). Scale bar, 1000 μm. OFR was defined as the number of organoids (≥50 μm) divided by the total seeded cells (n = 6). Data are representative of three independent experiments and indicated as means ± SDs. One-way analysis of variance (C and E) and two-tailed Student’s t-test (H) were utilized for statistics. \**P* <0.05, \*\**P* <0.01, \*\*\*\**P* <0.0001. NT, nontreated

Because only basal cells are capable of forming organoids (with terminally differentiated cells incapable of forming organoids), we used organoid formation to confirm if in the setting of FOXM1 inhibition there is decreased OFR due to enhanced differentiation. Indeed, both RCM-1 and siFOXM1 decreased the OFR which was further reduced by IL-13 treatment (Figure 3G and H and Supplemental Figure S1D and E).

### FOXM1 inhibition by RCM-1 enhances epithelial barrier integrity

Impaired epithelial differentiation disturbs the barrier integrity, which is critical in the pathogenesis of EoE^8,33^. In order to evaluate the ability of RCM-1 to restore epithelial barrier function, EPC2-hTERT cells were grown in ALI culture and given IL-13 to disrupt the epithelial barrier. Transepithelial resistance was significantly increased in cultures treated with RCM-1 despite IL-13-induced disruption (Figure 4A). IL-13-stimulated ALI cultures treated with RCM-1 demonstrated increased *IVL* and *FLG* with a decrease in *TP63* and *SOX2* (Figure 4B). Immunohistochemistry and immunofluorescence of ALI cultures similarly showed an increase in IVL and FLG, demonstrating increased differentiation after RCM-1 treatment, with a reciprocal decrease in TP63 (Figure 4C). These findings suggest that RCM-1 ameliorates impaired squamous stratification and barrier disruption by IL-13.

**Figure 4.**
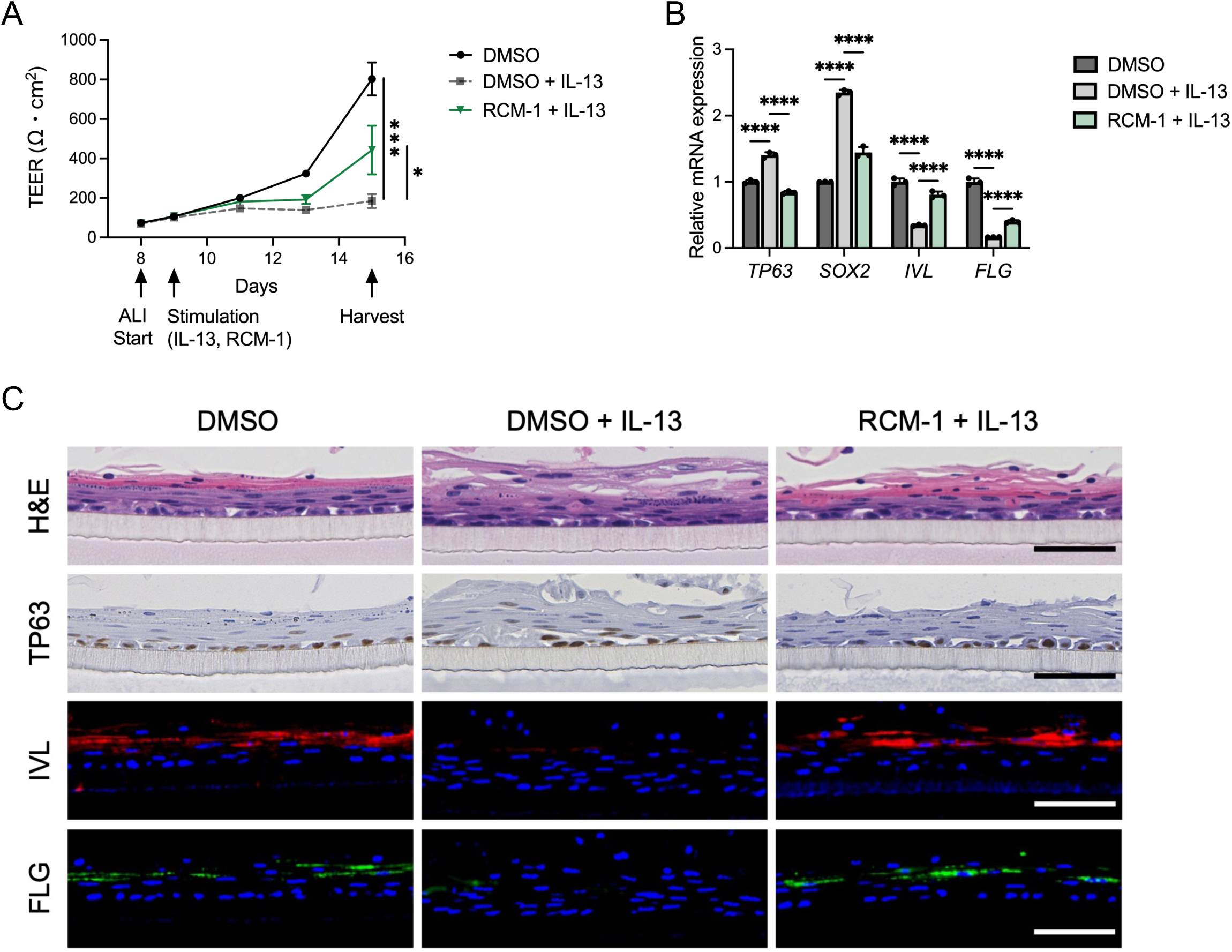
FOXM1 inhibition restores epithelial barrier integrity disruptions caused by IL-13 stimulation. (A) Transepithelial electrical resistance (TEER) of the EPC2-hTERT ALI-cultures (n = 3). EPC2-hTERT cells were kept in low-calcium (0.09 mM Ca^2+^) KSFM for 3 days, followed by high-calcium KSFM (1.8 mM Ca^2+^) for 5 days, and then remove media at day 8. Air-liquid interface (ALI)-cultured cells were stimulated with or without IL-13 (10 ng/ml) and RCM-1 (20 μM) from day 9 to 15. (B) Quantitative RT-PCR for *TP63*, *SOX2*, *IVL*, and *FLG* of the day 15 EPC2-hTERT ALI cultures (n = 3). (C) Representative images of hematoxylin and eosin (H&E), immunohistochemistry for TP63, and immunofluorescence staining for IVL (red), and FLG (green) of the day 15 EPC2-hTERT ALI-cultures. DAPI (blue). Scale bar, 50 μm. Data are representative of two independent experiments and indicated as means ± SDs. One-way analysis of variance (A and B) was utilized for statistics. \**P* <0.05, \*\*\**P* <0.001, \*\*\*\**P* <0.0001

### RCM-1 treatment improves epithelial changes and inflammation in murine EoE

Because FOXM1 inhibition with RCM-1 reversed the epithelial changes seen with IL-13 in vitro, we sought to determine the effects of RCM-1 in a murine model of EoE. RCM-1 administration reduced FOXM1 expression in mice undergoing the EoE protocol as well as untreated mice. In EoE-induced mice, RCM-1 decreased basal cell hyperplasia to levels comparable to non-EoE mice. Furthermore, RCM-1 treatment normalized the number of TP63 and Ki-67 positive cells in EoE mice (Figure 5B and C).

**Figure 5.**
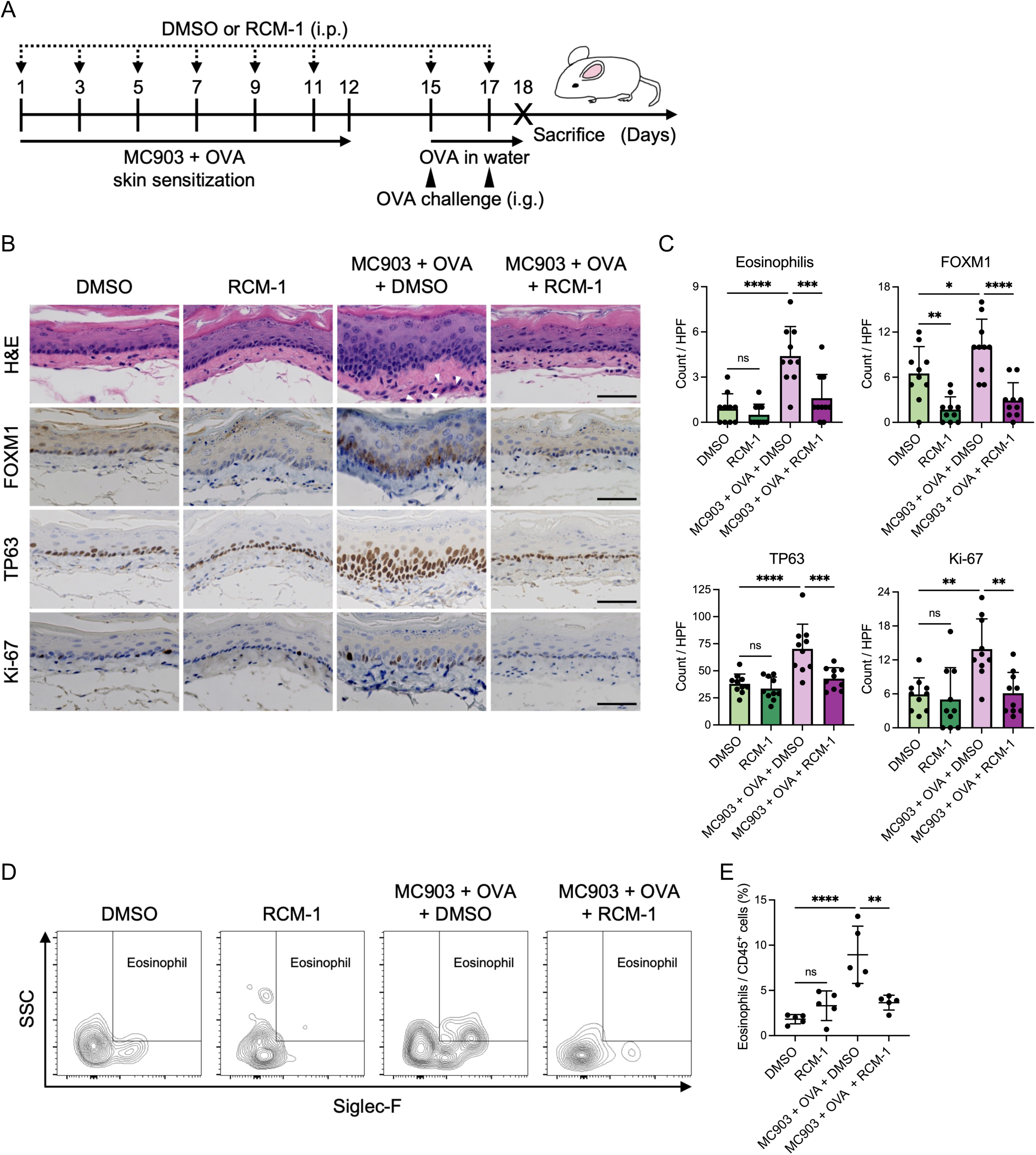
FOXM1 inhibition leads to decreases epithelial disruption and reduces inflammation in murine EoE model. (A) Schematic of murine EoE model and RCM-1 treatment. (B) Representative images of hematoxylin and eosin (H&E), immunohistochemistry for FOXM1, TP63, and Ki-67 of the murine esophagi. Arrowheads show infiltrating eosinophils. Scale bar, 50 μm. (C) Number of eosinophils per high-power field (hpf) in the esophagus. Number of FOXM1, TP63, or Ki-67-stained cells in the epithelium per hpf (n = 10). (D) Representative flow cytometry images of eosinophils in the murine esophagi. (E) Frequencies of eosinophils divided by CD45^+^ cells in the murine esophagi as measured by flow cytometry (n = 5). Data are indicated as means ± SDs. One-way analysis of variance (C and E) was utilized for statistics. \**P* <0.05, \*\**P* <0.01, \*\*\**P* <0.001, \*\*\*\**P* <0.0001. OVA, ovalbumin; i.p., intraperitoneal injection; i.g., intragastric gavage administration; ns, not significant

In addition to changes in the epithelium, there was decreased eosinophil infiltrate in EoE-induced mice treated with RCM-1 (Figure 5B-E), suggesting that FOXM1 has an effect on eosinophil chemotaxis. To investigate this finding further, we evaluated the effect of RCM-1 on *CCL26* expression (a major chemoattractant in the EoE esophagus)^6^. We found that RCM-1 treatment suppressed IL-13-induced *CCL26* expression in EPC2-hTERT organoids and PDOs (Figure 6A). Similarly, in biopsies of EoE patients, eosinophil count and *CCL26* expression were significantly and positively correlated with *FOXM1* expression (Figure 6B and C) Taken together, these suggest that FOXM1 inhibition improved the epithelial changes in EoE in addition to decreasing eosinophil chemotaxis.

**Figure 6.**
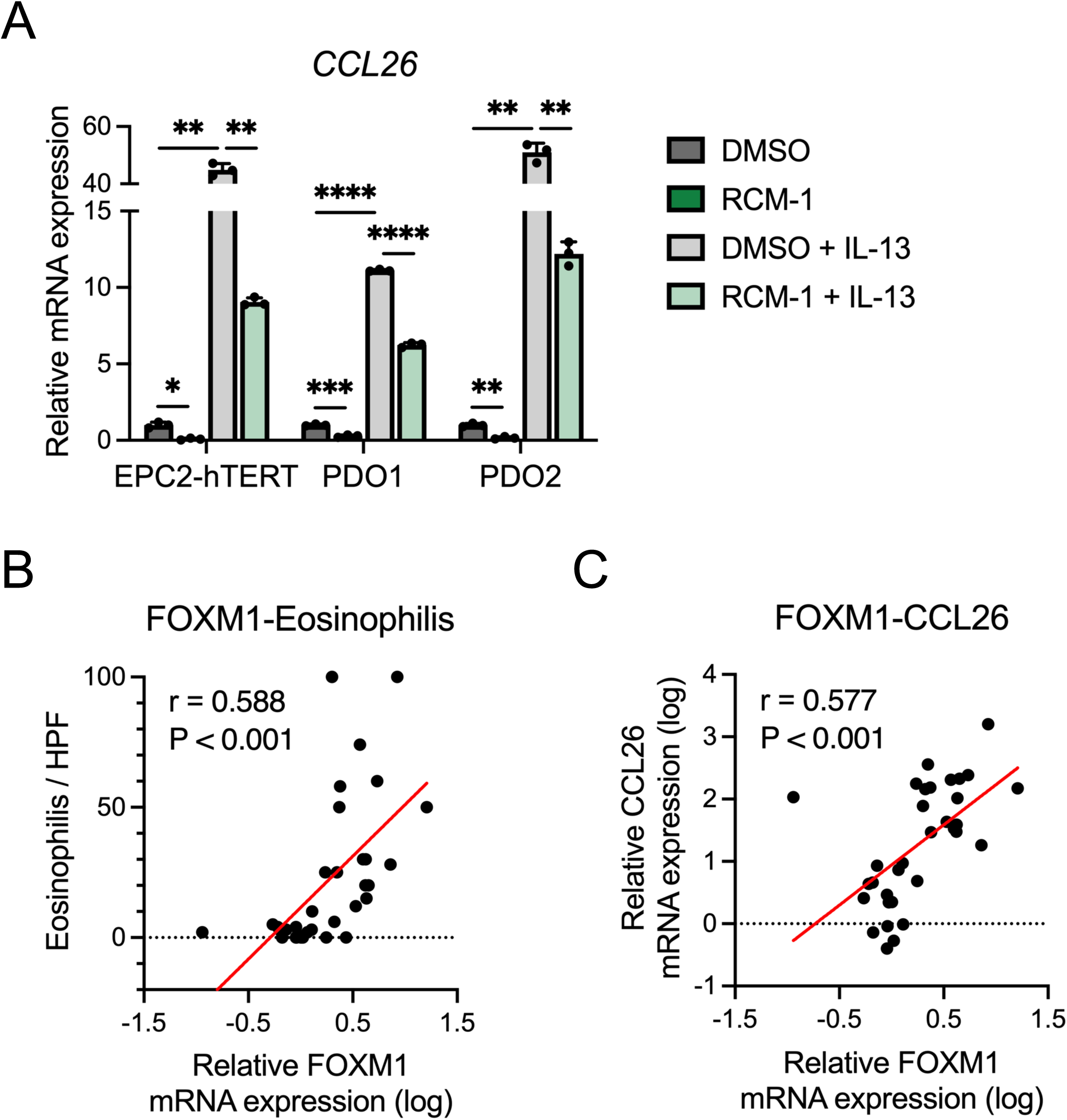
RCM-1 suppresses eosinophil chemotaxis in EoE. (A) Quantitative RT-PCR for *CCL26*. EPC2-hTERT or patient derived organoids (PDOs) were cultured for 7 days and then stimulated with or without IL-13 (10 ng/ml) and RCM-1 (10 μM) for 4 days. Day 11 organoids were harvested (n = 3). Data are representative of three independent experiments and indicated as means ± SDs. (B and C) Correlation analyses between *FOXM1* and eosinophils and *CCL26* on esophageal biopsies (n = 34). *FOXM1* and *CCL26* mRNA expressions were measured by quantitative RT-PCR and number of eosinophils per high-power field (hpf) were evaluated in the biopsies. One-way analysis of variance (A) and Pearson correlation coefficient (B and C) were utilized for statistics. \**P* <0.05, \*\**P* <0.01, \*\*\*\**P* <0.0001

### FOXM1 directly drives cell cycle and cellular proliferation in the esophageal epithelium

In order to understand the molecular changes that occur with reduction in FOXM1, we performed RNA sequencing of EPC2-hTERT monolayer cultures treated with siRNA FOXM1 and scrambled control. Gene ontology analysis revealed genes associated with cell adhesion were upregulated in the siFOXM1 cultures compared to siNC. On the other hand, genes associated with cell cycle were downregulated (Figure 7A). We then compared the DEGs from siFOXM1 cultures to the previously published EoE DEGs^25^. There was a marked reversal of expression genes found in the EoE transcriptome with FOXM1 knockdown (Figure 7B). Based on these findings, we sought to interrogate cell cycle within biopsies and siFOXM1 cells. In patient biopsies^25^, cell cycle gene CCNB1 was increased in EoE (Figure 7B and C). IL-13 stimulation also upregulated CCNB1 expression, while in the setting of FOXM1 knockdown, this gene was significantly downregulated (Figure 7D and E). Cell cycle analysis by WST-1 assay demonstrated decreased proliferative activity in the knockdown as well as a decrease in percent of cells in G2/M phase, contrary to changes by IL-13 stimulation (Figure 7F and G). Finally, we performed ChIP assay of FOXM1 and found that there was increased FOXM1 bound upstream near the transcription start site of CCNB1 (Figure 7H), suggesting that FOXM1 restores proliferation-differentiation gradient via transcriptional regulation of CCNB1 in EoE (Figure 7I).

**Figure 7.**
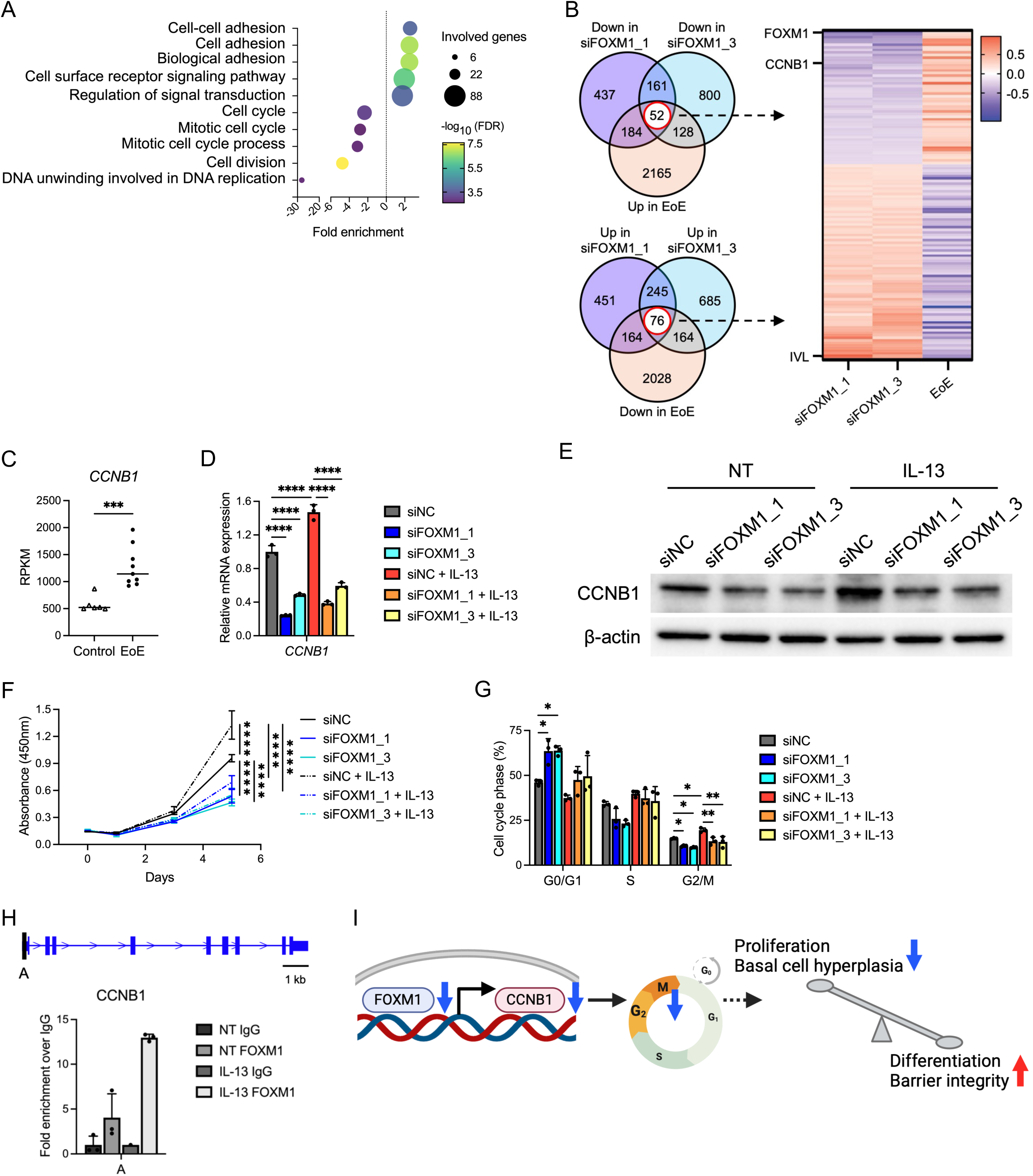
FOXM1 alters cell cycle progression via transcriptional regulation of CCNB1 in esophageal epithelium. (A) Top 5 enriched and depleted terms on differentially expressed genes (DEGs) overlapping in siFOXM1_1 and siFOXM1_3 cells based on Gene Ontology analysis. After the transfection, EPC2-hTERT cells were cultured in high-calcium (1.8 mM Ca^2+^) KSFM for 3 days and submitted to RNA sequencing. (B) Venn diagrams and heatmap generated by DEGs overlapping in the siFOXM1 and the EoE biopsy RNA sequencing (GSE58640). (C) Reads per kilobase per million (RPKM) values for CCNB1 in the EoE biopsy RNA sequencing (GSE58640) (control: n= 6, active EoE: n = 9). (D and E) Quantitative RT-PCR (n = 3) and representative images of immunoblot for FOXM1 in siFOXM1-transfected EPC2-hTERT cells in monolayer culture. The day following the transfection, cells were cultured in high-calcium (1.8 mM Ca^2+^) KSFM for 2 days along with or without 1 day of IL-13 (10 ng/ml) stimulation. (F) WST-1 assay for siFOXM1-transfected EPC2-hTERT cells (n = 3) (G) Cell cycle assay for siFOXM1-transfected EPC2-hTERT cells as measured by flow cytometry (n = 3). The day following the transfection, cells were cultured in high-calcium KSFM (1.8 mM Ca^2+^) with or without IL-13 (10 ng/ml) for 2 days. (H) Schematic of FOXM1 binding site identified on the promoter of CCNB1 and ChIP-qPCR for the binding site. Results were represented by fold enrichment method against IgG (n = 3; two samples of IL-13 IgG were not detected by qRT-PCR). (I) Schematic of the mechanism by which FOXM1 regulates the epithelial proliferation-differentiation gradient in esophagus. Data are representative of three independent experiments and indicated as means ± SDs. One-way analysis of variance (D, F, and G) and two-tailed Student’s t-test (C) were utilized for statistics. \**P* <0.05, \*\**P* <0.01, \*\*\**P* <0.001, \*\*\*\**P* <0.0001. FDR, false discovery rate; NC, negative control; NT, nontreated

### PI3K-AKT activation by IL-13 induces FOXM1 expression

While FOXM1 is known to be induced by the PI3K-AKT pathway in malignancy^34,35^, how it may be induced in the context of the Th2 inflammatory milieu of the EoE epithelium is unknown. We found increased phosphorylated PI3K in biopsies from patients with active EoE compared to non-EoE and inactive EoE (Figure 8A and B). Additionally, IL-13 induced phosphorylation of PI3K and AKT in EPC2-hTERT monolayer culture (Figure 8C). Treatment with PI3K/AKT pathway inhibitor LY294002 led to decreased expression of FOXM1 in EPC2-hTERT cells (Figure 8D). Together, our findings suggest a noncanonical role for IL-13 in stimulation of PI3K-AKT to drive FOXM1 expression in EoE (Figure 8E).

**Figure 8:**
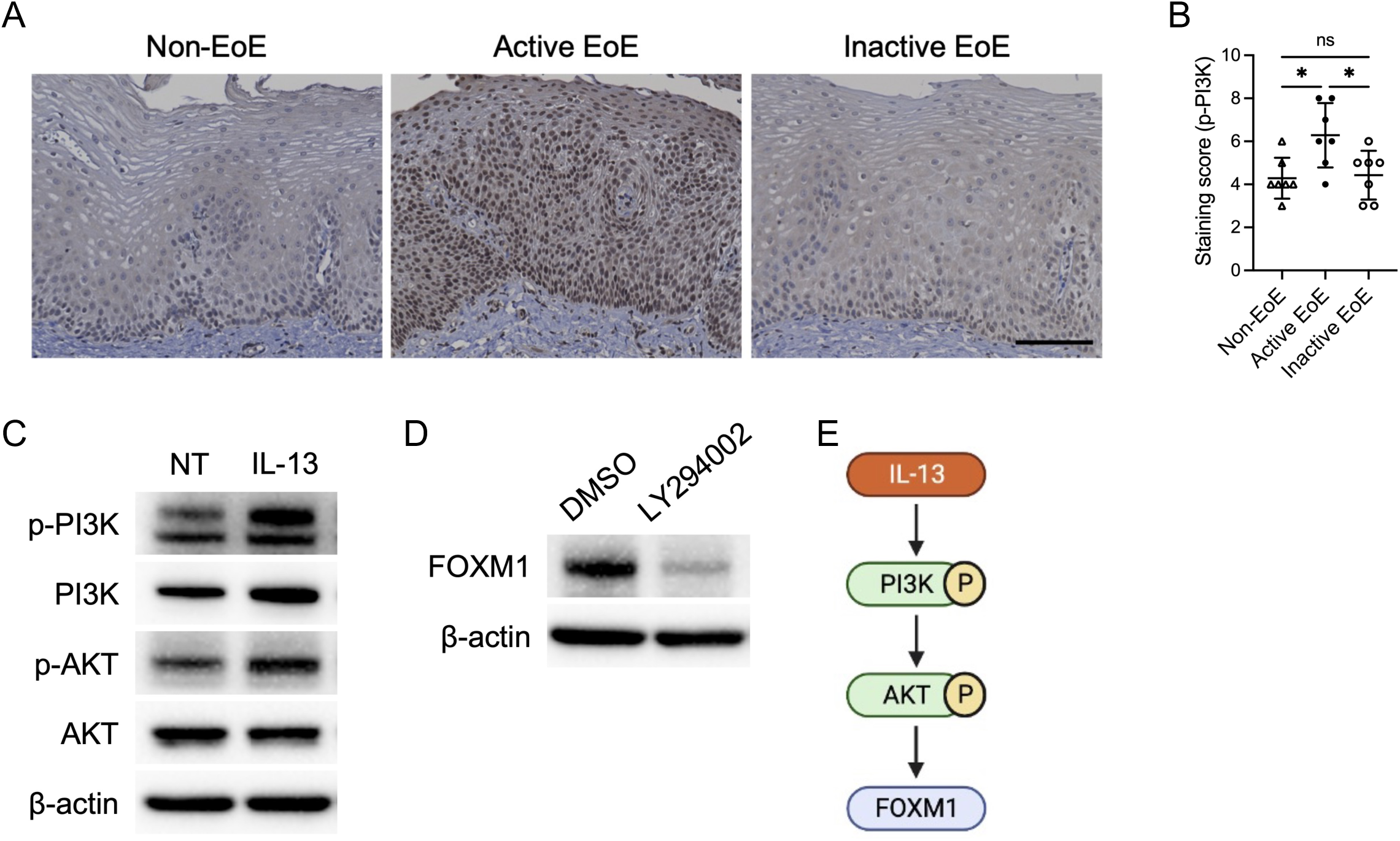
PI3K-AKT activation drives FOXM1 expression in EoE. (A and B) Representative immunohistochemical images of phospho-PI3K (p-PI3K) in biopsies from patients with non-EoE, active EoE, and inactive EoE. Scale bar, 100 μm. In the epithelium per high-power field, p-PI3K levels were quantified along with the following immunohistochemical score. Stained-nucleic intensity and stained-cytoplasmic area were score as 1, 2, 3, or 4, and then each value was added (n = 7 per group). Data are indicated as means ± SDs. (C) Representative images of immunoblot for p-PI3K, PI3K, p-AKT, and AKT in EPC2-hTERT cells stimulated with IL-13 (10ng/mL) for 30 min. (D) Representative images of immunoblot for FOXM1 in EPC2-hTERT cells treated with LY294002 (10μM) for 24 h. (E) Schematic of upstream mechanism of FOXM1 in esophageal epithelium. One-way analysis of variance (B) was utilized for statistics. \**P* <0.05. NT, nontreated; ns, not significant

## Discussion

Epithelial disturbances in EoE result from chronic inflammation and lead to diminished barrier integrity which normally protects the submucosa from insults such as food particles and gastric acid^36^. In order to identify promising therapeutic targets for EoE, it is imperative to elucidate the biological processes that regulate epithelial homeostasis. Herein we demonstrate that IL-13-induced differentiation defects and barrier disruption can be reversed by FOXM1 inhibition. We evaluate the therapeutic potential of FOXM1 inhibition demonstrating healing of the epithelium as well as decreased eosinophil chemotaxis. Mechanistically, we find that FOXM1 transcriptionally regulates proliferation in the esophageal epithelium and suggest a new role for PI3K/AKT pathway activation, driving cell proliferation through FOXM1 in EoE.

Previous studies have shown that Th2 cytokines promote esophageal epithelium proliferation in EoE^6,21,37,38^. In depth single-cell sequencing has shown an expansion of the proliferating suprabasal cells^39^. In conjunction with these hyperproliferating basal cells, the EoE epithelium is marked by a failure to differentiate^4^. We have previously shown that basal cell hyperplasia in the EoE esophagus associates with symptomatology^3^. Furthermore, in experimental models the epithelial disruption leads to barrier defects allowing for ongoing exposure to antigens^7,8,40^. However, the underlying mechanisms driving disruption in the proliferation/differentiation gradient in EoE are poorly delineated. Analyzing RNA sequencing data (GSE58640) of esophageal biopsies^25^, we demonstrated that cell cycle progression-related pathways, including PLK1, AURORA B, FOXM1, and AURORA A pathways, were notably enriched in patients with active EoE. FOXM1 is a transcription factor recognized as a master regulator of cell cycle and mitotic cell division^10,11^. FOXM1 is also crucial for embryonic development and progression of malignant tumors. Overexpression of FOXM1 has been found in a wide range of cancers, such as head and neck, esophagus, lung, and breast^41^, where it is involved in carcinogenesis, tumor growth, angiogenesis, and metastasis, contributing to poor prognosis^20,42,43^. Beyond its role in cell cycle and oncogenic processes, FOXM1 has recently been reported to be involved in epithelial cell differentiation in human skin keratinocytes^12–15^ and pulmonary allergic inflammatory disease contributing to goblet cell hyperplasia in asthma models^17,18^. Moreover, skin models demonstrate that FOXM1 upregulation induced expansion of the epithelial progenitor compartment with activated proliferative ability, which disturbed terminal differentiation, as seen in lack of keratin 13 and FLG, in keratinocytes, resulting in hyperplasia^12,13^. Conversely, depleted FOXM1 reduced proliferation and ameliorated the attenuated differentiation markers^14,15^. Taken together, these reports suggest that FOXM1 overexpression may result in hyperproliferation, leading to a defective terminal differentiation and barrier function.

To gain insight into the underlying mechanism of FOXM1 in esophageal epithelium, we performed RNA sequencing and ChIP analyses. As expected, cell cycle-related terms were decreased in siFOXM1-transfected EPC2-hTERT cells. FOXM1 accelerates entry into S and M-phase of cell cycle by transcriptionally regulating genes crucial for G1/S progression and G2/M transition, such as CCNB1, CDK1, CCNA2, and PLK1, promoting cellular proliferation^16,44–46^. We found that CCNB1 in particular was increased in EoE subjects and in vitro with IL-13 treatment; additionally, reduced by FOXM1 silencing. Of note, our ChIP experiments confirmed the binding of FOXM1 to the promoter region within 500 bp of CCNB1 transcription start site in esophageal epithelial cells. These findings suggest that FOXM1 directly regulates epithelial cell proliferation by modulating the G2/M transition via transcriptional regulation of CCNB1, with FOXM1 inhibition restoring the terminal differentiation and barrier function that are compromised by Th2 cytokines in EoE. This highlights a critical role for FOXM1 in the pathophysiology of EoE, where its modulation could potentially offer a strategy for restoring epithelial differentiation.

One unexpected finding from this work is that FOXM1 expression is also positively associated with the severity of EoE inflammation, specifically eosinophil counts in biopsies. In murine models of asthma, FOXM1 deletion reduced expression of CCL11 and CCL24 which in turn lead to decreased eosinophil and macrophage recruitment to the lung^18^, and RCM-1 treatment reduced the concentration of proinflammatory cytokines, such as IL-4, IL-5, IL-13, and IL-33^17^. Similarly, FOXM1 overexpression in the liver drove liver inflammation by directly binding to the promoter region of CCL2^47^. The fact that FOXM1 has dual roles, promoting epithelial disfunction and driving inflammatory infiltrate, makes this a particularly attractive therapeutic target. Our previous work as well as the work of others highlight the durability of epithelial changes in the EoE epithelium, with ongoing transcriptomic changes despite decreased eosinophil count^4,48^. Conversely, depletion of eosinophils in the esophagus does not lead to endoscopic or histologic changes in EoE^2^. A bidirectional approach, with medications targeting both the epithelium and immune compartments may offer more relief from inflammation, especially in refractory patients.

We sought to identify key upstream effectors of FOXM1 expression in the context of EoE. The PI3K/AKT signaling pathway is a major intracellular signaling pathway with aberrant activation identified widely in both malignant and non-malignant diseases. This pathway is critical for regulating numerous cellular processes, including proliferation, apoptosis, angiogenesis, migration, inflammation, and fibrosis^49,50^. Notably, beyond the canonical Janus kinase/STAT pathway, IL-13 has been shown to activate a non-canonical PI3K/AKT cascade in allergic diseases such as asthma and atopic dermatitis^51–53^. In cancer, this same PI3K/AKT axis plays a pivotal role in driving FOXM1 activity^34,35^. While it is established that PI3K/AKT signaling is essential for maintaining skin keratinocyte homeostasis^49^, its role within the esophageal epithelium in EoE remains largely unexplored. Our data reveal that phosphorylation of PI3K is significantly elevated in active EoE compared to both inactive EoE and control subjects. Moreover, we show that IL-13 directly activates PI3K and AKT, and that inhibition of the PI3K/AKT pathway using LY294002 effectively blocks FOXM1 expression in EPC2-hTERT cells. These findings underscore the critical role of IL-13 in driving FOXM1 upregulation via the PI3K/AKT signaling axis in the esophageal epithelium. As such, targeting the dysregulated PI3K/AKT network may offer novel therapeutic avenues for addressing the pathophysiology of EoE.

One limitation of our work is that while organoid and ALI culture are powerful 3D modeling techniques to simulate in vivo, these cultures consist of only esophageal epithelial cells, preventing the investigation of influences arising from interactions of epithelial cells with the heterogeneous cellular environment, such as fibroblasts, immune cells, and endothelial cells. Our recapitulated EoE milieu in vitro consists of IL-13, and there is concern that this does not take into account all inflammatory cytokine components and interactions. Co-culture with non-epithelial cells and other cytokines would be ideal to assess the role of FOXM1 in EoE in future work. The use of a murine EoE model ameliorates some of these concerns and shows feasibility of RCM-1 treatment. However, an artificially induced and non-human model cannot completely recapitulate a human disease.

Taken together, we highlight the role of FOXM1 as a key modulator in the balance between the epithelial proliferation and terminal differentiation in the esophagus. FOXM1 inhibition may restore epithelial homeostasis and barrier integrity to mitigate hyperproliferation and basal cell hyperplasia through regulation of cell cycle-related gene and inflammatory cytokine in response to allergic inflammation, leading to improved symptoms in patients with EoE. Thus, approaches directly targeting the epithelial transcriptional network for the homeostasis may represent a promising therapeutic strategy for EoE treatment.

## Acknowledgments

We thank to the Molecular Pathology and Imaging Core (Kate Bennett, Rebecca Ly, and Jonathan P Katz) for technical support. Schematics were created with BioRender.com.

## Data availability statement

The data and materials described in this study are available upon reasonable request to the correspondence.

## Ethics Statements

### Patient consent for publication

Not applicable.

## Ethics approval

This study was approved by the Institutional Review Board of the Children’s Hospital of Philadelphia (no. 10-007737) and the Hospital of the University of Pennsylvania (no. 813841). After obtaining informed consent, human esophageal tissues and clinical information were collected from the participants. All animal experiments were performed in accordance with the Institutional Animal Care and Use Committee of The Children’s Hospital of Philadelphia.

