## Supplementary figures and images for "FOXM1 Modulation Alleviates Epithelial Remodeling and Inflammation in Eosinophilic Esophagitis"

### Supplemental Figure 1

Supplemental figure S1

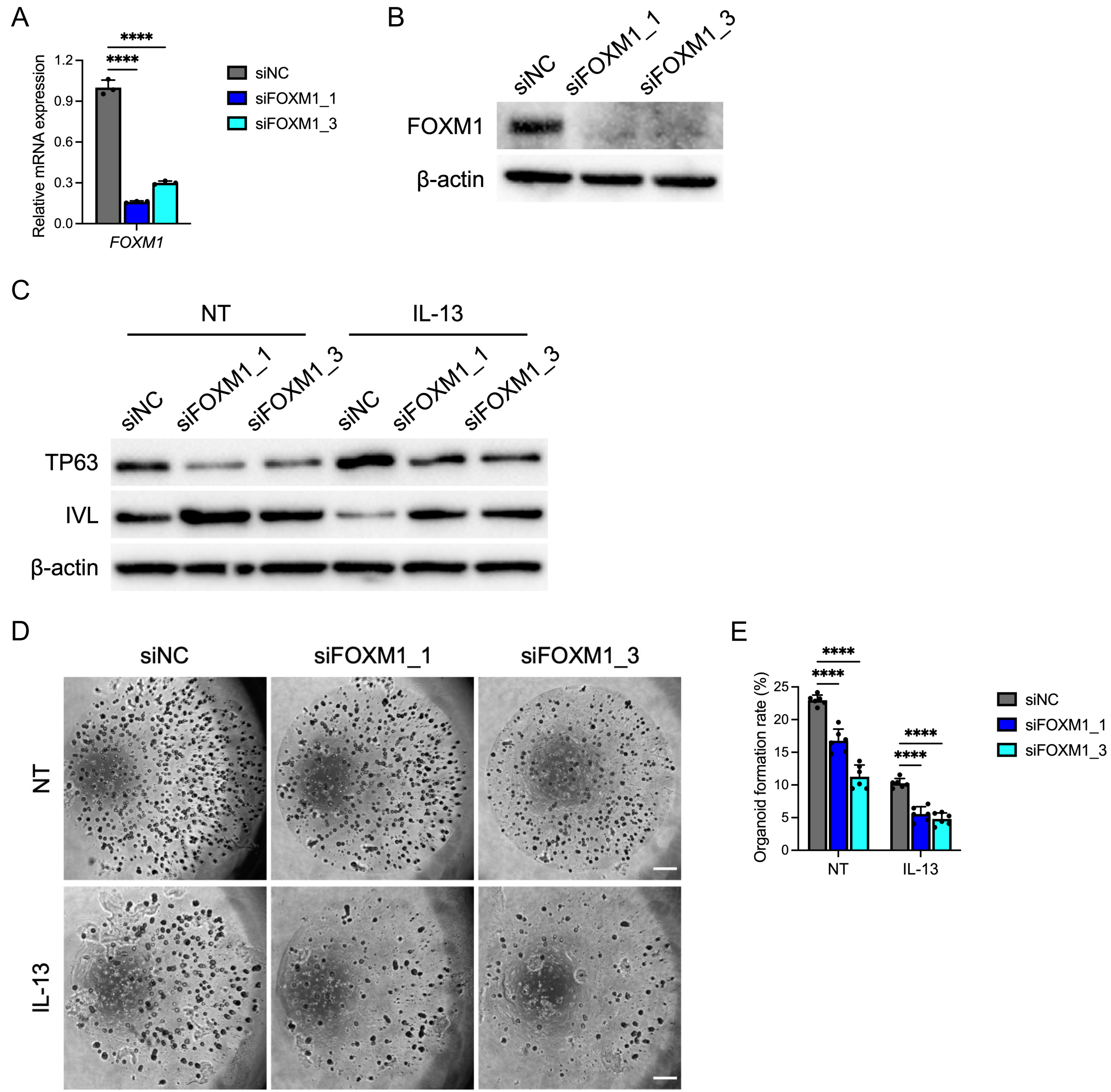

### Supplemental Figure 2

Supplemental figure S2

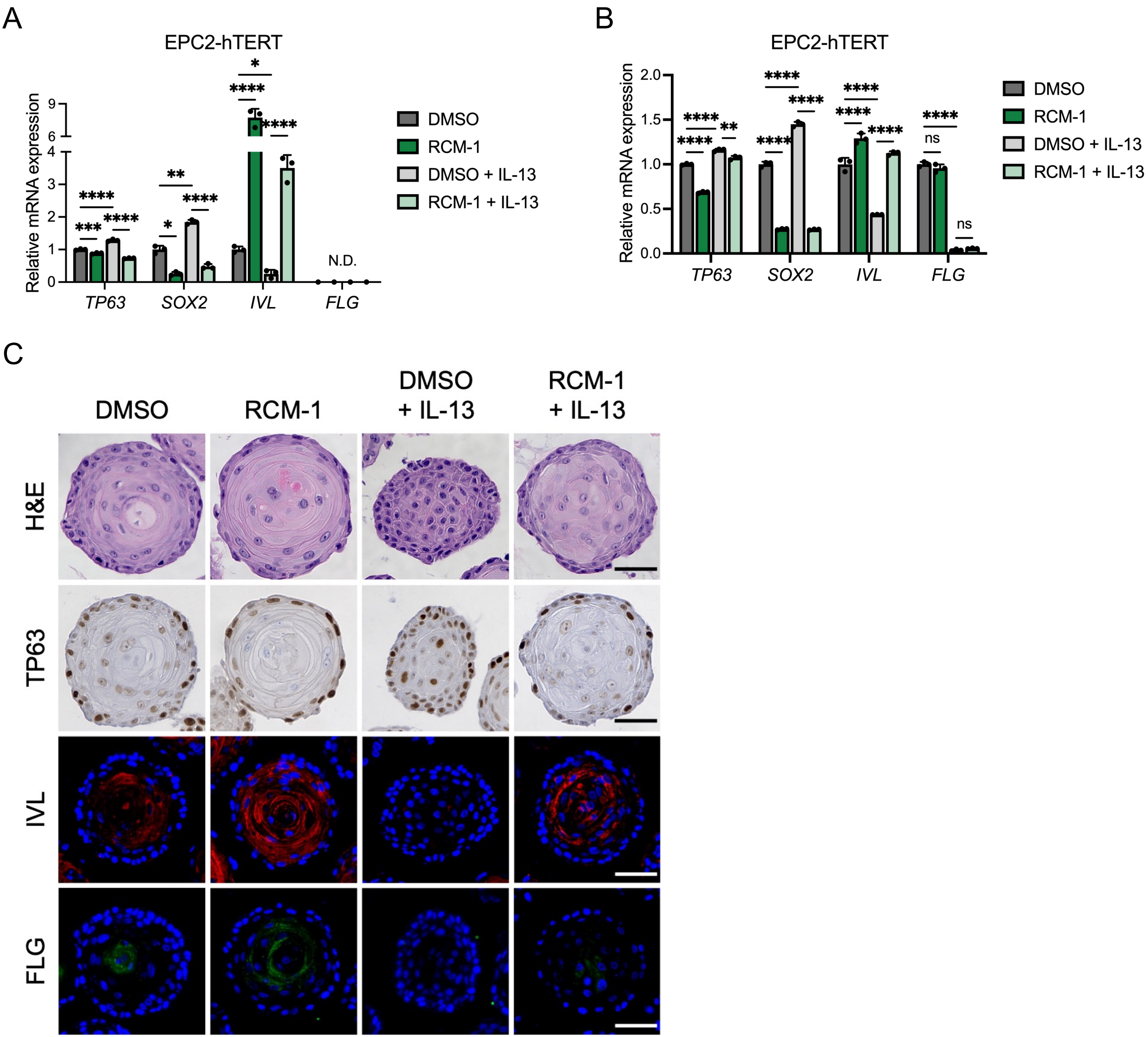
