## Supplementary material for "FOXM1 Modulation Alleviates Epithelial Remodeling and Inflammation in Eosinophilic Esophagitis": Supp Methods

**Supplemental information.**

**Methods**:

*Monolayer culture of human esophageal cells*

Immortalized human normal esophageal epithelial cell line EPC2-hTERT cells were kept in keratinocyte-serum free medium with 0.09 mM Ca^2+^ (KSFM; Thermo Fisher Scientific, Waltham, MA) containing human recombinant epidermal growth factor (1 ng/ml), bovine pituitary extract (50 μg/ml), and 1% penicillin-streptomycin. High-calcium (1.8 mM Ca^2+^) KSFM was used to induce differentiation. The cells were cultured in a 5% humidified CO_2_ incubator at 37°C^1–3^. Cells were stimulated by recombinant human IL-13 (213-ILB; R&D Systems, Minneapolis, MN), IL-4 (204-IL; R&D Systems), tumor necrosis factor alpha (TNF-α; 210-TA; R&D Systems), or transforming growth factor beta (TGF-β; 240-B; R&D Systems) to recapitulate EoE milieu. The Robert Costa Memorial drug-1 (RCM-1; S6898; Selleck Chemicals, Houston, TX) and LY294002 (S1105; Selleck Chemicals) was used to inhibit FOXM1 activity and phosphatidylinositol 3-kinase (PI3K) signaling, respectively^4,5^.

*Esophageal 3-dimensional organoid culture*

EPC2-hTERT or primary patient derived organoids (PDOs) were established as described previously^6–8^. Briefly, human esophageal biopsies were minced in dispase (10 U/ml; Corning Inc., Corning, NY) and incubated for 10 min at room temperature. After removing dispase, the biopsies were kept in 0.05% trypsin and mixed with a thermomixer at 700-800 rpm for 10 min at 37°C. Dissociated pieces were filtered through a 70 μM cell strainer and collected in soybean trypsin inhibitor (250 mg/ml; Sigma-Aldrich, St. Louis, MO). The cells were forced through a 35 μM cell strainer again and resuspended in KSFM. Following dissociation into a single cell suspension, EPC2-hTERT and the patient derived cells were suspended in Matrigel basement membrane matrix (Corning Inc.) and seeded under modified KSFM with 0.6 mM Ca^2+^. Y27632 (10 μM; Sigma-Aldrich) was added for patient derived organoids only on the first day. Organoids were cultured for 7 days and then treated with IL-13 (10 ng/ml), RCM-1 (10 μM), or vehicle (phosphate-buffered saline for IL-13 or dimethyl sulfoxide (DMSO) for RCM-1) for 4 days. The day 11 organoids were harvested for further analyses.

Organoid formation rate (OFR) was defined as the number of ≥50 μm organoids at day 11 divided by the total seeded cells in each well^6,9^.

*Air-liquid interface (ALI) culture system*

EPC2-hTERT cells were cultured on transwell permeable supports (3470; Corning Inc.) with KSFM containing 0.09 mM Ca^2+^ for 3 days. After reaching confluence, media were changed to KSFM with 1.8 mM Ca^2+^. In order to induce epithelial stratification and differentiation, the media in the apical component was removed at day 8. Cells were stimulated by adding IL-13 (10 ng/ml), RCM-1 (20 μM), or vehicle in the basolateral compartment from day 9 to 15. Transepithelial electrical resistance (TEER) was measured by Epithelial Volt/Ohm (TEER) Meter (World Precision Instruments, Sarasota, FL) to evaluate epithelial barrier. The ALI-cultured epithelium at day 15 was harvested for RNA extraction and histology.

*Patients and esophageal specimens*

After obtaining informed consent, distal esophageal specimens were derived from patients who underwent esophagogastroduodenoscopy (EGD). Based on diagnostic criteria of EoE^10^, we classified patients with the histologic presence of ≥15 esophageal mucosal eosinophils (eos) per high-powered field (hpf) as active EoE, patients with EoE who resolved histologic inflammation and eosinophilia (<15 eos/hpf) upon follow-up EGD as inactive EoE, and patients who had no previous diagnosis of EoE and complained of symptoms justifying EGD but demonstrated no endoscopic or histopathologic abnormalities as non-EoE. Patients with a history of inflammatory bowel disease, gastrointestinal bleeding, celiac disease, or any other acute or chronic intestinal disorders were removed from recruitment^11^. This study was approved by the Institutional Review Board of the Children’s Hospital of Philadelphia (no. 10-007737) and the Hospital of the University of Pennsylvania (no. 813841). Patient demographics are shown in Supplemental Table S1. Patient and Public Involvement were not appropriate or possible to involve patients or the public in the design, or conduct, or reporting, or dissemination plans of our research.

*Murine EoE models*

EoE models were induced as described previously^8,12,13^. Briefly, female BALB/c mice (The Jackson Laboratory, Bar Harbor, ME) were sensitized epicutaneously with 2 nmol MC903 (Tocris Bioscience, Bristol, UK) in ethanol and 1000 μg ovalbumin (OVA; A5503-50G; Sigma-Aldrich) for 12 days, followed by intragastrical challenge with 50 mg OVA on days 15 and 17. Drinking water was switched to 15 g/l OVA-containing water on day 15. Mice were treated intraperitoneally with 3 mg/kg RCM-1 or vehicle, prepared with 5% DMSO and 95% corn oil, every other day. Mice were euthanized on day 18. Esophagi were harvested for flow cytometry (5 mice per group) and histology (10 hpf views randomly selected from 3 mice per group). All experiments were performed in accordance with the Institutional Animal Care and Use Committee of The Children’s Hospital of Philadelphia.

*Quantitative reverse transcription-polymerase chain reaction (qRT-PCR)*

RNA isolation and reverse transcription were performed as described previously^1,11,14^. Real-time qRT-PCR was carried out with TaqMan Gene Expression Assays (Thermo Fisher Scientific) or Fast SYBR Green Assays (Thermo Fisher Scientific) using the StepOnePlus Real-Time PCR System (Thermo Fisher Scientific). The used TaqMan and SYBR Green primers are provided in Supplemental Table S2 and S3. Relative mRNA levels of each gene were standardized against *glyceraldehyde-3-phosphate dehydrogenase* (*GAPDH*) levels as a housekeeping control.

*Western Blot (WB)*

Whole-cell lysates were extracted as described previously^9^. Equivalent amounts of protein (20-40 µg) were subjected to a NuPAGE 4 to 12% Bis-Tris gel, electrophoresed, and then transferred to a polyvinylidene difluoride membrane. After 1 h blocking with 5% non-fat dry milk or bovine serum albumin, membranes were incubated with primary antibodies overnight at 4°C. Primary antibodies used are provided in Supplemental Table S4. A horseradish peroxidase (HRP)-conjugated secondary antibody (1:2000; NA934 or NA931; Amersham BioSciences, Buckinghamshire, UK) and Clarity Western ECL Substrate (Bio-Rad Laboratories, Hercules, CA) were used to detect immunoblots. β-actin was set as a loading control.

*Immunohistochemistry and immunofluorescence*

Paraffine-embedded human and murine esophagi, organoids, and ALI-cultured epithelia were sectioned and subjected to hematoxylin and eosin (H&E) staining, immunohistochemistry, or immunofluorescence as described previously^9,15^. Briefly, for immunohistochemistry, sections were stained with primary antibodies. The signal was found with VECTASTAIN Elite ABC-Peroxidase Kit (Vector Laboratories, Burlingame, CA). Diaminobenzidine was utilized for color modification. For immunofluorescence, sections were incubated with primary antibodies at 4°C overnight. 4′,6-diamidino-2-phenylindole (DAPI; 17985-50; Electron Microscopy Sciences, Hatfield, PA) was utilized to stain nuclei. Stained slides were imaged with an All-in-One Fluorescence Microscope BZ-X710 (KEYENCE Corp., Osaka, Japan). Primary antibodies used are provided in Supplemental Table S4.

*Flow cytometry*

Murine esophagi were dissociated and filtered in the same manner as human specimens described above. The single-cell droplets were resuspended in phosphate-buffered saline containing 1% bovine serum albumin. Dead cells were detected by LIVE/DEAD Fixable Aqua Dead Cell Stain Kit (L34957; Life Technologies, Carlsbad, CA). Viable cells were incubated with anti-CD11b (101243; BioLegend, San Diego, CA), anti-CD45 (564279; BD Biosciences, Franklin Lakes, NJ), and anti-Siglec F antibody (565527; BD Biosciences). Eosinophils were identified as CD45^+^/CD11b^+^/Siglec F^+^/SSC^high^ cells^16^. BD LSRFortessa Cell Analyzer (BD Biosciences) and FlowJo software (FlowJo LLC, Ashland, OR) were used for analyses.

*Cell cycle assay*

The day after the transfection, the media were switched to the high-calcium KSFM with or without IL-13 (10 ng/ml) for 2 days. Cells were then fixed with 70% ethanol for 2 days, rinsed, and stained with propidium iodide. Samples were evaluated through BD LSRII (BD Biosciences). Cell cycle phases were provided by the Watson Pragmatic algorithm in FlowJo software^17^.

*Chromatin Immunoprecipitation (ChIP) assay*

ChIP samples were prepared with truChIP Chromatin Shearing Kit (520154; Covaris, Woburn, MA) based on the manufacturer's protocol. Briefly, 15 million EPC2-hTERT cells were harvested in a 15 cm dish and fixed with 11.1% formaldehyde for 10 min for cross-linking. After quenching, nuclei were collected and transferred into Millitube 1 ml AFA Fiber (520135; Covaris). Prepared chromatin was shared by Covaris S2 Focused Ultrasonicator (Covaris). Sonication setting was as follows; duty cycle 5, intensity 4, cycles/burst 200, time 8 min. Shared chromatin was diluted to 1:1 with IP Dilution Buffer (in the kit) and then centrifuged. Immunoprecipitation was performed with SimpleChIP Plus Sonication Chromatin IP Kit. The supernatant was transferred into a microcentrifuge tube. After removing the diluted chromatin for input, 5 μg of FOXM1 antibody (sc-376471 X; Santa Cruz Biotechnology, Dallas, TX) or IgG (2729; Cell Signaling Technology) was added and incubated overnight at 4℃ with rotation. Lysis was then incubated with 30 μl of magnetic beads (9006S; Cell Signaling Technology, Danvers, MA) for 2 h 15 min at 4℃ with rotation. After low and high salt wash, chromatin was incubated for 30 min at 65℃ to elute from the antibody and beads and then 2 h at 65℃ for reverse crosslinking. Finally, DNA was purified using QIAquick PCR Purification Kit (28104; QIAGEN, Hilden, Germany) and quantified by real-time qPCR with SYBR Green technology according to the manufacture’s protocol. ChIP results were calculated by fold enrichment method against IgG. The primer against cyclin B1 (CCNB1) for ChIP-qPCR is provided in Supplemental Table S3.
