## Supplementary material for "FOXM1 Modulation Alleviates Epithelial Remodeling and Inflammation in Eosinophilic Esophagitis": Supp Table 1

**Supplemental Table S1. Patient demographics in this study**

| **ID** | **Application** | **Age (years)** | **Gender** | **Diagnosis** | **Eos/hpf** | **History of impaction** | **History of stricture** | **Symptom of dysphagia** | **Symptom of regurgitation** |
| --- | --- | --- | --- | --- | --- | --- | --- | --- | --- |
| 1 | PCR | 7.3 | Male | Non-EoE | 0 | No | No | No | Yes |
| 2 | PCR | 11.3 | Male | Non-EoE | 0 | No | No | No | Yes |
| 3 | PCR | 7.2 | Male | Non-EoE | 0 | No | No | No | Yes |
| 4 | PCR | 14.5 | Male | Non-EoE | 0 | No | No | No | No |
| 5 | PCR | 6.3 | Male | Non-EoE | 0 | No | No | No | Yes |
| 6 | PCR | 13.8 | Female | Non-EoE | 0 | No | No | Yes | Yes |
| 7 | PCR | 9.9 | Male | Non-EoE | 0 | No | No | No | Yes |
| 8 | PCR | 53.0 | Female | Non-EoE | 0 | No | No | Yes | No |
| 9 | PCR | 45.0 | Female | Non-EoE | 0 | No | No | Yes | No |
| 10 | PCR | 27.0 | Male | Non-EoE | 0 | No | No | No | No |
| 11 | PCR | 49.0 | Female | Non-EoE | 0 | No | No | No | No |
| 12 | PCR | 51.0 | Male | Non-EoE | 0 | Yes | No | Yes | No |
| 13 | PCR | 36.0 | Female | Non-EoE | 0 | No | No | Yes | Yes |
| 14 | PCR | 13.3 | Female | Active EoE | 100 | Yes | No | No | No |
| 15 | PCR | 9.3 | Male | Active EoE | 20 | No | No | Unknown | Unknown |
| 16 | PCR | 10.2 | Male | Active EoE | 25 | No | No | No | No |
| 17 | PCR | 12.0 | Male | Active EoE | 28 | Yes | No | No | Yes |
| 18 | PCR | 7.8 | Male | Active EoE | 100 | No | No | No | No |
| 19 | PCR | 9.6 | Male | Active EoE | 60 | Yes | No | No | No |
| 20 | PCR | 9.3 | Male | Active EoE | 15 | No | No | No | No |
| 21 | PCR | 11.2 | Male | Active EoE | 30 | Yes | No | No | No |
| 22 | PCR | 10.0 | Male | Active EoE | 50 | No | No | Yes | Yes |
| 23 | PCR | 17.3 | Female | Active EoE | 25 | No | No | Yes | No |
| 24 | PCR | 48.0 | Male | Active EoE | 20 | Yes | No | No | No |
| 25 | PCR | 24.0 | Female | Active EoE | 30 | No | No | Yes | Yes |
| 26 | PCR | 28.0 | Female | Active EoE | 50 | Yes | No | No | Yes |
| 27 | PCR | 45.0 | Female | Active EoE | 58 | Yes | Yes | No | Yes |
| 28 | PCR | 25.0 | Male | Active EoE | 74 | No | Yes | Yes | Yes |
| 29 | PCR | 3.3 | Male | Inactive EoE | 0 | No | No | No | Yes |
| 30 | PCR | 16.2 | Female | Inactive EoE | 0 | No | No | No | No |
| 31 | PCR | 9.9 | Male | Inactive EoE | 1 | No | No | No | No |
| 32 | PCR | 50.0 | Female | Inactive EoE | 2 | Yes | No | Yes | No |
| 33 | PCR | 45.0 | Male | Inactive EoE | 4 | Yes | No | Yes | No |
| 34 | PCR | 47.0 | Male | Inactive EoE | 0 | Yes | No | No | No |
| 35 | PCR | 49.0 | Male | Inactive EoE | 5 | Yes | No | No | No |
| 36 | PCR | 18.0 | Male | Inactive EoE | 0 | Yes | No | No | No |
| 15b | PCR | 9.0 | Male | Inactive EoE | 2 | No | No | No | No |
| 16b | PCR | 9.3 | Male | Inactive EoE | 6 | No | No | No | Yes |
| 17b | PCR | 12.3 | Male | Inactive EoE | 3 | Yes | No | Unknown | Unknown |
| 18b | PCR | 7.5 | Male | Inactive EoE | 3 | No | No | No | No |
| 19b | PCR | 9.0 | Male | Inactive EoE | 12 | Yes | No | No | No |
| 20b | PCR | 8.6 | Male | Inactive EoE | 0 | No | No | No | No |
| 22b | PCR | 9.5 | Male | Inactive EoE | 10 | No | No | Yes | No |
| 25b | PCR | 24.0 | Female | Inactive EoE | 2 | No | No | Yes | No |
| 26b | PCR | 29.0 | Female | Inactive EoE | 4 | Unknown | Unknown | Yes | No |
| 28b | PCR | 25.0 | Male | Inactive EoE | 0 | No | Yes | Unknown | Unknown |
| 30b | PCR | 17.2 | Female | Inactive EoE | 0 | No | No | No | No |
| 37 | IHC | 16.1 | Male | Non-EoE | 0 | Yes | No | No | No |
| 38 | IHC | 16.8 | Female | Non-EoE | 0 | Yes | No | No | No |
| 39 | IHC | 17.3 | Female | Non-EoE | 0 | Yes | No | Yes | Yes |
| 40 | IHC | 15.7 | Female | Non-EoE | 3 | Yes | No | Yes | Yes |
| 41 | IHC | 14.9 | Male | Non-EoE | 0 | Yes | No | Yes | No |
| 42 | IHC | 5.0 | Male | Non-EoE | 0 | No | No | No | No |
| 43 | IHC | 14.0 | Female | Non-EoE | 0 | Yes | No | Yes | Yes |
| 44 | IHC | 3.9 | Female | Active EoE | 80 | No | No | Unknown | Unknown |
| 45 | IHC | 18.3 | Male | Active EoE | 60 | No | No | Unknown | Unknown |
| 46 | IHC | 5.4 | Female | Active EoE | 30 | No | No | Yes | No |
| 47 | IHC | 15.3 | Male | Active EoE | 60 | Yes | No | Unknown | Unknown |
| 48 | IHC | 12.9 | Male | Active EoE | 32 | No | No | Unknown | Unknown |
| 49 | IHC | 1.3 | Male | Active EoE | 15 | No | No | Unknown | Unknown |
| 50 | IHC | 7.4 | Female | Active EoE | 15 | No | No | Yes | Yes |
| 51 | IHC | 14.1 | Female | Inactive EoE | 0 | No | No | No | No |
| 52 | IHC | 14.7 | Male | Inactive EoE | 3 | Yes | No | No | No |
| 53 | IHC | 8.9 | Male | Inactive EoE | 0 | No | Yes | Yes | Yes |
| 54 | IHC | 3.5 | Male | Inactive EoE | 1 | No | No | No | No |
| 55 | IHC | 4.2 | Female | Inactive EoE | 0 | Yes | No | No | No |
| 56 | IHC | 12.7 | Male | Inactive EoE | 0 | No | No | No | No |
| 57 | IHC | 14.7 | Male | Inactive EoE | 10 | No | No | No | No |

Eos, eosinophil; HPF, high-power field; PCR, polymerase chain reaction; IHC, immunohistochemistry
