## Supplementary material for "FOXM1 Modulation Alleviates Epithelial Remodeling and Inflammation in Eosinophilic Esophagitis": Supp Table 2

**Supplemental Table S2. List of TaqMan probes used for qRT-PCR**

| **TaqMan probe** | **Catalog number** |
| --- | --- |
| FOXM1 | Hs01073586_m1 |
| SOX2 | Hs01053049_s1 |
| TP63 | Hs00978340_m1 |
| IVL | Hs00846307_s1 |
| FLG | Hs00856927_g1 |
| CCL26 | Hs00171146_m1 |
| GAPDH | Hs02786624_g1 |
